# Ecological Representations

**DOI:** 10.1101/058925

**Authors:** Sabrina Golonka, Andrew D Wilson

## Abstract

Mainstream cognitive science and neuroscience both rely heavily on the notion of representation in order to explain the full range of our behavioral repertoire. The relevant feature of representation is its ability to *designate* (stand in for) spatially or temporally distant properties, When we organize our behavior with respect to mental or neural representations, we are (in principle) organizing our behavior with respect to the property it designates. While representational theories are a potentially a powerful foundation for a good cognitive theory, problems such as grounding and system-detectable error remain unsolved. For these and other reasons, ecological explanations reject the need for representations and do not treat the nervous system as doing any mediating work. However, this has left us without a straight-forward vocabulary to engage with so-called ‘representation-hungry’ problems or the role of the nervous system in cognition.

In an effort to develop such a vocabulary, here we show that James J Gibson’s ecological information functions to designate the ecologically-scaled dynamical world to an organism. We then show that this designation analysis of information leads to an ecological conceptualization of the neural activity caused by information, which in turn we argue can together support intentional behavior with respect to spatially and temporally distal properties. Problems such as grounding and error detection are solved via law-based specification. This analysis extends the ecological framework into the realm of ‘representation-hungry’ problems, making it as powerful a potential basis for theories of behavior as traditional cognitive approaches. The resulting analysis does, according to some definitions, allow information and the neural activity to be conceptualized as representations; however, the key work is done by information and the analysis remains true to Gibson’s ecological ontology.

## Introduction

Mainstream accounts of cognition, and the role of the brain in cognition, are predicated on theories of (computational) mental and neural representation. These cognitive representations are not invoked on a whim; they take the forms they do because they are meant to enable three important features of cognition, 1) flexibility / intentionality, 2) successful perception, 3) higher order cognition. However, a number of issues with cognitive representations remain; the neural code by which they are instantiated in the brain is unknown, there is no broad agreement on their structure and format, and they may be fatally ungrounded (e.g. Harnad, 1990) and shut off from system detectable error (Bickhard, 2009). As long as cognitive neuroscience relies on cognitive representational theories, it inherits these problems. In the absence of alternatives, this might be an acceptable risk. If, on the other hand, a framework existed that could explain the key features of cognition while avoiding the problems that saddle cognitive representations, then neuroscience would be on firmer footing by adopting this theoretical basis instead.

This paper attempts specifically to show how an alternative, ecological framework can explain all three key features of cognition that motivated early cognitive theorists while avoiding the problems of grounding and system-detectable error. We note that Gibsonian ecological information solves the poverty of stimulus problem via specification, and we then show that the details of this solution means information *designates* (stands in for) behaviorally relevant properties of the currently present world to cognitive systems. We then propose that this information interacts with nervous systems in a way that allows the resulting neural activity to serve as components that can designate more spatially and temporally distal properties for the cognitive system, and so support flexible, intentional, ‘higher-order’ cognition. We also argue that, because ecological information is a real, physically identifiable entity, with a lawful relationship to designated properties, it is a more grounded foundation for neuroscience than cognitive representations. Consequently, we think that this is a project worth pursuing.

A brief note: this issue of designation connects directly to the notion of representation (hence the name of the paper), which obviously trips alarm bells for ecological psychologists. We will return to this point at the end.

## Why Cognitive Science Invokes Representations

A good theory should explain the characteristics of a set of related phenomena. A good cognitive theory needs to explain the full range of behaviors we can perform, and most of these behaviors go beyond mechanical cause-and-effect. It’s relatively uncomplicated to tell a causal story about simple, mechanically linked events. If we are sitting in a tree and the branch we are sitting on breaks, the force of gravity causes us to be displaced. The branch temporarily overcame the force of gravity which pulled us towards the ground. When this oppositional force was removed, our mass accelerated towards the ground according to the laws of motion. It’s more challenging when events are not so obviously mechanically linked. For instance, if we move because a branch falls nearby, or because it is windy and we are worried that a branch might fall, what is it that causes our behavior to change? In this example, there was no mechanical linkage between the accelerating tree branch and our body; in the latter case, there wasn’t even an accelerating tree branch at all, just the worry of one. Yet, we moved. Obviously, we don’t move by accident or by magic. There are reasons and explanations, but these reside in a psychological level of explanation, which is why we need a psychological theory to fill the gap.

Newell (1980) proposed a list of characteristics that a theory of cognition needs to be capable of addressing. To him, the most important feature is universality, which is the idea that a cognitive system must be able to behave as an (almost) arbitrary function of the environment. It must be able to be ‘about’ anything it encounters in the world; cognition must be *intentional*. Newell emphasized that implementing such a flexible, adaptive system in an actual, physical system was a major problem that must be addressed (Bechtel, 1998; Stich, 1992).

We can divide the intentionality requirement into two sub-cases:

2. *Intentionality with respect to spatially distal properties.* We can act intentionally towards properties of objects/events that aren’t in direct mechanical contact with us, via our perceptual contact with those objects/events. Cognitive theories consider perception to be impoverished and ambiguous, meaning that successful intentional interaction with the environment requires a system to supplement/structure perceptual contact with stored knowledge. But, even non cognitive approaches must explain how organisms coordinate behavior with respect to environmental properties that aren’t in direct mechanical contact.
3. *Intentionality with respect to higher order or temporally distal properties.* We can act intentionally towards higher-order properties of objects/events that might not be present in, or might even be in opposition to, the current environment. Here, our perceptual contact with such entities is non-existent, so again our experience must be supplemented to support successful intentional interactions.

Early cognitive scientists (e.g. Fodor, 1980; Pylyshyn, 1989) argued that universality could only be implemented in a *computational* system because these have the requisite indefinite flexibility and can in principle be realized in a variety of physical systems. It is commonly argued that computational systems are necessarily representational^1^. Therefore, the primary motivation for cognitive psychology to treat cognition as necessarily representational is that computation has been the best way to explain how cognitive intentionality and flexibility could be (at least in theory) implemented in a physical system. The current mainstream consensus is therefore that cognition requires *mental* representations that are *internal* to the system (specifically, implemented in the brain^2^) and *computational* in nature.

How do these representations gain traction on the problem of intentionality? Ramsey (2007) has analyzed the concept of representation in extensive detail. He has identified that in order to have any bite, representation has to mean something more than simply an “inner” or “causally relevant” state (p 8). While there remain many arguments about the format and content of cognitive representations, Ramsey argues that, regardless of bells and whistles, all cognitive theories of representation rest upon “the basic idea that inner structures in some way serve to stand for, *designate*, or mean something else” (Ramsay, p 3, emphasis ours). Newell (1980) usefully defines designation like this:

> Designation: An entity X designates an entity Y relative to a process P, if, when P takes X as input, its behavior depends on Y.
>
> There are two keys to this definition: First, the concept is grounded in the behavior of a process. Thus, the implications of designation will depend on the nature of this process. Second, there is action at a distance … This is the symbolic aspect, that having X (the symbol) is tantamount to having Y (the thing designated) for the purposes of process P (Newell, 1980, p. 156).

Cognitive representations therefore try to fulfill the intentional job description of a good theory of cognition on the basis of their computational, representational ability to designate otherwise inaccessible properties to a cognitive system.

### Problems with Cognitive Representations

Though cognitive representations remain central to cognitive models, they have not been accepted uncritically, even from within the pro-representation camp. There is no consensus on the format and structure of cognitive representations, nor how representations are instantiated in neural activity. Worse, there are still critical concerns about whether representations, of any structure or format, can do the intentional work cognitive scientists need them to do.

The first of these concerns is that representational systems (of any type) do not necessarily come with intrinsic access to intentional content and, thus, cannot actually be ‘about’ anything without external help (the *symbol grounding problem*^3^, Bickhard, 2009; ‘empty symbols’, Harnad, 1990; Searle, 1980). Symbolic representations, for example, allow an arbitrary mapping between the structure of the representation and the thing it’s meant to represent. This provides crucial flexibility for universality, but because the mapping is arbitrary there’s no guaranteed way for the representation to have any content, let alone the right content (this problem has most recently been laid out in detail in the interface theory of perception; Hoffman, Singh & Prakesh, 2015; see also Wilson, 2018a).

Attempts to solve the grounding problem often try to link abstract or higher-order mental representations to more basic perceptual representations (e.g. Barsalou, 1999; Harnad, 1990). Such well-motivated attempts fail, however, due to the lack of an adequate theory of perception (Turvey et al, 1981; Wilson, 2018a) which means that even the lower perceptual representations lack internally defined content. This “infinite regress of interpreters interpreting” (Bickhard, 2009, p 573) is endemic to any representational account where content is defined externally, which Bickhard argues includes all the main types of representation (including the theories of Milikan, Fodor, Dretske, and Cummins).

The second, related concern is that symbolic/representational systems do not automatically come with a frame of reference to identify when they are making errors. Without a way to identify when a system is behaving incorrectly, behavior cannot adapt to become better attuned to the environment (Bickhard, 2009). Error-guided learning requires that a system be able to detect representational errors. But, this ability depends on access to representational content – a system can’t know that it’s wrong if it’s blindly trading in symbols with inaccessible content. An example is the arbitrary relationship between letters and sounds. If you have ever tried to help a child learn to read^4^, you will know that you cannot simply hand them a piece of paper with words on; there is nothing intrinsic to the representational letter system that allows the child to know what the sound is or when they are making a mistake. Learning to read is possible, but detecting an error requires extensive support from outside the representational system, in the form of a teacher, and even then it is extraordinarily difficult to learn. Given the absence of an external teacher, the relationship between cognitive representations and the things they represent must be internally available within the bounds of the representational system in order for system-detectable error to be possible. According to Bickhard’s analysis, no current representational accounts achieve this.

For these and other reasons, non-representational theories of cognition are becoming more common. These are, for the most part, derived from Gibson’s ecological approach to perception-action (Gibson, 1979).

### Ecological Alternatives

#### Affordance-Based Approaches

To get around these problems, a variety of ‘radical’ theories of cognition argue that one or more of the key features of cognition can be successfully explained without invoking representations (e.g. Chemero, 2009; Gibson, 1979; Kono, 2009; Reitveld & Kiverstein, 2014; Schmidt, 2007). These theories typically rest on Gibson’s demonstration that there is no poverty of stimulus. There is now evidence for a wealth of high-quality perceptual information available to support coordination with spatially distal properties, and so the case for representations is undercut in this behavioral domain.

To try and move into the domain of more ‘representation-hungry’ problems (higher order and/or temporally distal properties), these approaches expand Gibson’s notion of affordances to include any relation that can be defined between an organism and its environment, up to and including social, linguistic, and cultural relations (Chemero, 2009; Reitveld et al, 2014). Unfortunately, these theories lack a mechanism by which these relational affordances might be perceived. If an organism cannot come into psychological contact with such relational affordances, then they are not a suitable basis for a theory of cognition (Golonka, 2015; Wilson, 2018b). For this reason, we argue that any attempt to scale up ecological psychology must begin with the ecological information organisms are in contact with, and how organisms can use information in the selection and control of increasingly complex and abstract behaviors.

#### An Information-Based Approach

The proposed components of a cognitive theory must support intentional behavior, including cases where the behaviorally-relevant properties of objects or events are spatially and/or temporally distal. Furthermore, the components must have grounded intentional content and enable system-detectable error. Neither cognitive representational theories nor affordance-based non-representational theories can, to date, fulfill this entire job description. Cognitive psychology therefore still has a bill to pay, and because cognitive neuroscience is cognitive psychological theories, it is on the hook for the debt.

In order to successfully address all three key features of cognitive systems, we propose an expansion of the ecological approach grounded in the use of ecological information. We will argue that ecological information functions to designate the world to an organism. This framing emerges quite naturally from careful consideration of the nature of ecological information; how it comes to be, and how it is used by an organism. We then argue that information shapes behavior by changing the spatiotemporal activity of the nervous system so that this activity designates the information to the action systems. We then propose that the designation relationship between environmental properties and ecological information, and between information and the neural activity it creates, supports intentional behavior with respect to the behaviorally-relevant properties of objects or events, even those that are not currently present in the organism’s task environment. Thanks to specification, this chain of designation remains firmly grounded and capable of supporting error detection and correction. As a result, we suggest that this theoretical perspective is the best current starting point for understanding how the nervous system mediates between perception and action. The rest of this paper steps through this analysis in detail.

## An Ecological Approach to Cognition & Neuroscience

Our approach to a general ecological theory of cognition is grounded in the nature of ecological information. In this section, we will explain what information is, its function to the organism, and what it does to the nervous system. With this in hand, we will identify how these components fulfill the job description for a theory of cognition.

### Ecological Information Specifies Environmental Properties

Ecological information is constituted by higher-order relational patterns in energy arrays. These patterns are created by the lawful interaction of the energy array with the dynamics of the world (Turvey et al, 1981) and are used by organisms to perceive that world (Gibson, 1966, 1979; Michaels & Carello, 1981).

Properties of objects and events in the world change over space and time in ways that reflect the composition and organization of these properties. Consequently, these properties are best described at the level of *dynamics* (Bingham, 1988, 1995) which allows units of time, position (and its derivatives), and mass. Our perceptual systems are not in direct mechanical contact with most environmental properties of interest, however. Instead, they are in contact with energy media like light and sound. Structures in energy media reflect the composition and structure of the local environment. But, whereas properties of the environment are best described at the level of dynamics, structures in energy media are only kinematic projections of these properties (Bingham, 1988; Turvey et al, 1981). Kinematics is a level of description that only refers to motions; the units are time, position (and its derivatives), but *not* mass. For a given property, the kinematic description will not be identical to the dynamic description because mass is not represented in kinematics (the ‘perceptual bottleneck,’ Bingham, 1988).

Fortunately, properties of the environment are projected into energy media via a law-based process. The details of the projection are unambiguously related to the details of the dynamic that caused it, and different dynamical properties project differently. Consequently, the kinematic patterns *specify* (i.e., map 1:1 to) the dynamical properties that caused them, without needing to be identical to them (Gibson, 1979; Turvey et al, 1981). Ecological information is, therefore, a *kinematic specification of dynamical properties* (Runeson & Frykholm, 1983). Its function is to enable an organism to be connected to those properties without requiring mechanical contact so we can successfully interact with a (typically distant) task environment.

### Ecological Information Designates the Dynamical World to Organisms

When we talk about an organism ‘successfully interacting with a task’ it means that the organism’s behavior complements the dynamics of the task. From an ecological stance, a system that exhibits intentionality is one whose behavior is appropriately organized with respect to task dynamics.

Almost all behaviorally relevant dynamics in the world are ‘over there’; they are spatially or temporally distal and not in mechanical contact with the organism; and yet, all animals can couple and coordinate behavior with respect to some subset of relevant distal properties. We achieve this by being in contact with lawfully created, specifying kinematic projections of those dynamics into the energy media those dynamics are embedded in. We humans and all other animals are also embedded in these energy media, and we have evolved specialized perceptual systems that reliably react to contact with structured energy arrays^5^. Therefore, the projection of dynamic properties into energy arrays connects task dynamics to organisms. We suggest that it is useful to think of the function of information for organisms as designating environmental properties.

With reference to Newell’s (1980) definition described above, P’s behavior depends on the specified dynamical property Y; and the structure of P is only explainable with reference to X and its relationship to Y. The general finding in ecological behavioral research shows that the form of a behavior is coordinated and controlled with respect to task-relevant dynamical properties, but that the details of the behavior map tightly onto the structure of the information involved. For example, the behavioral-level characteristics of coordinated rhythmic movement are accounted for by the structure of relative direction, which is the specifying information for the dynamical property relative phase (Bingham, 2001, 2004a, b, reviewed in Golonka & Wilson, 2012, 2018; see Wilson et al, 2005 for a key empirical demonstration). This means that, while coordinated rhythmic movement is intentional with respect to a dynamic property of the world, organisms achieve coordination only by using a kinematic information variable as a stand-in. Importantly, as demonstrated by Wilson et al (2005), we can detect this distinction experimentally – it is not merely a theoretical stance. These results unambiguously support the claim that behavior (P) complements the environment (Y) by virtue of the organism’s contact with information (X).

The consequence of ecological information on perception-action systems is, therefore, best explained by invoking the ability of information to designate properties of objects and events in the world. In fact, we’d struggle to explain why organisms react to structure in kinematic arrays without invoking the ability of these patterns to designate world properties.

Usefully, this particular designation is firmly grounded and supports error detection and correction. Ecological information is the result of a law-based process, which makes the content of that information non-arbitrary; the consequent neural activity we discuss in the next section then inherits that grounding (Harnard, 1990). Errors *can* occur when the information variable being used does not specify the currently relevant task dynamics (because it is the wrong variable to use, or because of an experimental perturbation). However, because of specification, nothing additional to the information is required for it to mean what it does and be used to coordinate and control behavior; information is internally related to its contents (c.f. Bickhard, 2009). Any mismatch is therefore readily available to the organism and can drive corrections. There is a growing experimental literature on this type of error correction (e.g. Jacobs, Michaels & Runeson, 2000; Jacobs, Runeson & Michaels, 2001; van der Meer et al, 2012; Withagen & van Wermeskerken, 2009) which in turn has led to the development of the direct learning theory (Jacobs & Michaels, 2007). Information is functioning as a stand-in for the distal environment, but it happens to be an excellent one.

### Neural Activity Designates Information to Behavioral Systems

Property-specifying patterns in kinematic arrays provide a mechanism connecting the world to perceptual systems. We also need a mechanism connecting information to behavior. It is obvious that the nervous system is the primary connection between information and behavior, but our ecological analysis presents a decidedly novel way to characterize what the nervous system needs to accomplish. Specifically, we must explain how information causes changes in nervous system activity such that resultant actions complement environmental properties. Given that behavior typically tracks the form of the information being used, an initial parsimonious hypothesis is that the nervous system supports this by preserving ecological informational structure.

Explicitly ecological analyses of neural activity (i.e. tracking the neural consequences of interacting with ecological information) are currently rare, but the studies that do exist support our hypothesis (e.g. Agyei et al, 2015; Magrassi et al (2015); van der Meer et al, 2012; van der Weel & van der Meer, 2009). For example, van der Meer et al measured the magnitude of the correlation between looming-related information variables and neural activity. They found that the strength of this relationship predicted task performance. In fact, changes in the magnitude of the correlations in an individual predicted changes in performance over time. In other words, task performance (the extent to which behavior complements task dynamics) depended on how well informational structure was preserved in neural activity.

This reconceptualization of the neural code follows naturally from the ecological analysis of behavior. Rather than operating by synchronizing oscillations at various frequencies, the language of the brain may be more akin to the continuously unfolding nature of ecological information. The existence of this type of ongoing activity, similar to so-called travelling waves, has been known about for decades (see Hughes, 1995 for a review), but it has not made much of an impact on neuroscience. One reason is that common neuropsychological techniques (e.g., EEG recording at the scalp, averaging across trials) effectively obscure travelling-wave effects (Muller et al, 2018). In addition, there is no way to really understand what the structure of nervous system activity is doing without looking at the structure of energy entering the system. The ecological conception of information is the first coherent theory of how patterns in energy media relate to relevant environmental properties, and, therefore, is the best existing theoretical account from which to analyze how the spatiotemporal structure of neural activity enables intentional behavior.

### An Information-Based Theory of Cognition

We now have a set of ecological components (information and the neural activity it creates) that can enable us to propose a general ecological account of cognition and the brain.

Information immediately enables the first two key features of cognitive systems:

*1. Intentionality:* Because ecological information variables specify biologically relevant task dynamical properties (rather than perceptual primitives), they are inherently meaningful (Turvey et al, 1981) and support behavioral flexibility (Newell’s universality). Even a completely novel object or event will have *properties* in common with things we have already encountered, which will cause familiar kinematic projections and provide a basis for exploring the new context. In the absence of access to familiar properties, behavior falls apart (e.g., during a whiteout).
*2. Action with respect to spatially distal properties:* The lawful process by which the dynamical world is projected into information enables the kinematic specification of those dynamics. While there is a physical distance between us and most of the world, this distance is literally filled with structure in energy media that is specific to biologically relevant properties of distal objects and events. This law-based process also supports grounding and system-detectable error as the content is internally defined.

To tackle the third motivation (*action with respect to temporally distal properties*) we must now move beyond well-trodden paths for ecological psychology to build an argument about how information and consequent neural activity can support ‘higher order cognition’ and ‘thinking about things in their absence’.

## Ecological ‘Higher-Order’ Cognition

Research on ecological information focuses on the continuous control of action. This tells us a great deal about how information structures real-time, online interactions with the world. But, especially for humans, behavior often reflects the influence of things that are not currently present in the environment. For example, a person might choose to have a decaf coffee at lunch after remembering how much regular coffee they drank that morning. This moves us into the territory of so-called ‘representation-hungry’ problems (Clark & Toribio, 1994). Even Barsalou (1999), who worked very hard to defend the importance of perception in cognition, hypothesized that perceptual experience is reified into perceptual symbols, which could form the basis of “higher” cognitive functioning.

The ecological solution we develop below remains firmly grounded in ecological information. Our goal is to identify how and when some portion of spatiotemporal structure of neural activity caused by ecological information might become decoupled from the information. Decoupled activity provides a cognitive resource to organisms, which allows the possibility of structuring behavior with respect to temporally distal properties. We will now step through this in detail.

### Action Control, Action Selection, Neural Representations

Our developing solution begins by identifying that information can both control actions and *select*^6^ them (Golonka, 2015). Action selection occurs when an organism chooses between alternatives, changes from one task to another, or parameterizes the performance of the current task. When a friend asks you to ‘pick up the red cup’, information in the auditory signal enables you to select which of two cups you pick up. You then use visual and proprioceptive information specifying the location of the cup and the movement of your arm to implement and control the action.

The two roles (action control and action selection) place different requirements on information. In order for information to support action *control*, it must change in behaviorally-relevant real time as a direct function of some task-relevant property in the environment. In other words, it must continuously specify the current state of the world that created the information (e.g. as relative direction does for relative phase; Wilson & Bingham, 2008). This is what enables ecological information variables to support real-time coupling of behavior to properties currently present in the task environment.

But while all ecological information is lawfully related to the properties of objects or events in the world that create the information, organisms are *not* law-bound to use that information in a particular way (Wilson 2018b). Following Golonka (2015), when the behavioral consequences of information are not related to the object or event that caused the information, we say information has had a *conventional* (as opposed to law-based) effect on behavior.^7^

If one encounters a door that says “Danger: Bear Inside,” the behaviorally-relevant properties of the bear do not structure patterns in ambient light that reach your retina because the bear is occluded by a door. However, ecological information caused by the sign on that door can cause neural activity that participates in selecting actions related to these distal properties (e.g., avoidance). The fact that some scribbles on a sign shape functional action selection reflects a long-term immersion of a particular type of nervous system (i.e., human) in a particular type of learning context (i.e., one that encourages written communication). But, critically, from the first person perspective of the organism, it is just interacting with information.

For our purposes, the relevant distinction between information used to select versus control actions concerns the stability of consequent neural activity. There is no convincing evidence that we can decouple neural activity sufficient to support action control from the presence of the relevant information in the environment. One example of this that experienced drivers, despite being able to successfully steer a real car, are unable to realistically mime the action of steering, and often do so in ways that would have catastrophic consequences in actual driving contexts (Wallis et al, 2007). Without ongoing informational support, action control eventually falls apart. Knowing how to steer in a real driving context is not the same as, and does not entail, making our nervous system act as if it is steering absent that context. This basic distinction (in the form of the online vs offline control of actions) also shows up in most of the core work around the two visual streams hypothesis; see Goodale, et al, (2004) for a review.

In contrast, we *are* often able to instantiate neural activity corresponding the spatiotemporal structure of information used in action selection. This is likely to be the case if 1) we have an appropriate precipitating event and 2) the structure of the information is simple, short, and/or well-practiced and stereotypical enough to have had a reliable functional effect on corresponding neural activity during learning. In humans, a familiar example of such neural activity is the experience of inner speech. The structure of individual words for an experienced language user is simple, short, well-practiced and relatively stereotypical. The right precipitating event (e.g., reading the sign “Danger: Bear Inside”) can reliably instantiate neural activity with the spatiotemporal structure of the acoustic information caused by pronouncing these words. The result is that we “hear” the words in our heads (e.g. Breen & Clifton, 2013). This example relies on a close relationship between information present in the moment and the neural activity (i.e., they contain the same words in different modalities), but this connection is not obligatory. We could imagine training someone on a convention that a red circle on a door means that there is a bear inside^8^. In this case, the information created by the colored circle causes neural activity that is structurally similar to the auditory information caused by the word “bear.” This neural activity functions as a stand-in for the acoustic ecological information for the word “bear.”^9^ The structure of the activity has been shaped by the repeated presence of the acoustic information in real life. But now, the neural activity is stable enough to be instantiated by an appropriate precipitating event. Through this relationship, the neural activity corresponding to the spatiotemporal structure of the word “bear” can impact action selection – for example, by selecting avoidance behaviors. Once an action is selected, the actual escape from the bear will require access to online information suitable for action control; the neural activity corresponding to the word “bear” cannot tell your legs how to move with respect to the supporting surface of the floor.

This is a simple example of how ecological information can enable functional behavior with respect to things not in the present environment, via things that are. There are two things worth drawing attention to. First, neural activity decoupled from ecological information present in the environment only has the power to select actions, not to control them. Second, decoupled neural activity isn’t simply an example of mental representation grounded in perceptual experience. They are re-instantiations of neural activity caused by ecological information. Such re-instantiations can have a phenomenality; it feels like something to hear language in our heads or to imagine someone’s face; and they can have a consequence on future behavior by impacting selection of actions or selection of other neural activity (e.g., we continue our train of inner speech). But this neural activity, while internal, is not the same as a mental representation in standard cognitive theories.

An ecological framework that includes both designating information and the neural activity it creates can therefore provide a psychological level of analysis that connects intentional behavior to temporally distal environmental properties. Thus, this framework is robust enough to address all three key features of cognitive systems outlined earlier.

### How to Handle More Abstract Cognition

With this analysis in place, we will now discuss how neural instantiations of action-selecting information could enable some of the trickier aspects of human cognition. Only neural activity that meets our working definition of designation (i.e., that designates an environmental property with respect to corresponding ecological information) will be considered. Also, let us recall that not all such neural activity will be stable enough to instantiate in the absence of the corresponding information. In fact, there should be a distribution of stability, such that some neural representations can be re-instantiated quite accurately (in the sense of a strong systematic relationship between neural activity and information, rather than an exact replica of the structure of the information), some with a certain degree of accuracy, some with very poor accuracy, and some, not at all. For the purposes of this discussion, we are concerned with the subset of neural activity that can be re-instantiated with fair to good accuracy.

In order to support counterfactual and other higher-order cognition, cognitive scientists often argue that knowledge systems must be both *conceptual* and *componential*, to allow complex expressions to be decomposed and new expressions to be built up, i.e. *productivity* (Chomsky, 1957; Dietrich & Markman, 2003; Fodor & Pylyshyn, 1988; Haugeland, 1991). Conceptual systems are removed from the particulars of a situation – they can represent general cases (concepts) rather than individuals (Barsalou, 1999). Componential systems contain parts that can be combined (and re-combined) according to, for example, a recursive syntax. Symbol systems can quite clearly realize these features, which is what makes them such a good option for supporting counterfactual thinking and context-dependent flexibility^10^.

Counterfactual thinking is clearly a capacity of at least some cognitive systems. Thus, our main challenge is to show that an ecological approach can support counterfactual thinking. We don’t have any stock in whether or not this solution involves a conceptual and componential system, so if the ecological solution does not have these features, it isn’t really a blow to our primary argument. However, as it turns out, we think that certain features of our ecological components mean that they do enable conceptual, componential systems, making their ability to support higher-order cognition easier to see.

It is uncontroversial to say that developing a *concept* requires experience with multiple individuals of a type. For us, this would involve (at a minimum) repeated exposure to ecological information variables specifying properties of a given type. The neural activity caused by this repeated exposure will vary in many respects based on the details of the individuals and differences in neural states when information makes contact with the nervous system. However, if the individuals tend to share any ecologically specified properties (i.e., if they really are a type) then there will be correspondingly stable aspects of shared neural activity, invariants over the transformation of experience^11^. This subset of neural activity would acquire a certain degree of stability, such that the activity can be re-instantiated, given the right precipitating event, in the absence of the corresponding information. Because ecological information designates properties and not individuals, this kernel of stable neural activity can designate properties associated with a type. This means that ecological neural activity of a certain kind can function as concepts.

This same subset of neural activity can also support *componential* systems. We predict that stable neural activity will only emerge if the corresponding information is sufficiently simple, short, and stereotyped. This type of neural activity is a component – it is a bit that can participate in a number of events made up of other bits. These ecological neural components then enable *productivity* in the following way. Neural activity of this kind can impact action selection. Some of these actions can be the instantiation of further neural activity. Some of the variance in what actions are selected by a given neural activation will be explained by the learning history of the organism. For instance, if the acoustic event for the letter “B” almost always follows the acoustic event of the letter “A” in a person’s learning history, then activating “A” will tend to select the activity for “B” (such as when you start singing the ABC song in your head). But some of the variance will also be explained by the current context, summarized in the informational environment and current neural and bodily state of the organism. So, if you are watching West Side Story, then activating “A” may select the subsequent activity “…Jet is a Jet is a Jet all the way.” Therefore, ecological neural components can be combined in multiple ways with other neural components and the grounded way they do so is functionally related to learning history and current context^12^. In addition, because information designates properties, not individuals, it and the related neural activity don’t suffer from the holism that, some argue (e.g. Barsalou, 1999) makes typical perceptual theories unable to support componentiality.

Conceptual, componential, and productive systems support aspects of higher-order cognition like counterfactual thinking, thinking about impossible things, and talking about imaginary things. As we said before, we believe that the important task for us here is to show how ecological psychology can support higher-order cognition, whether or not the ecological solution also requires a conceptual and componential system. But, if the reader endorses the logic that aspects of higher-order cognition naturally follow from concepts, componentiality, and productivity, then we hope to have shown how ecological neural activity possess these features. We think this demonstration does some important work in justifying the viability of our approach, but we would like to add one final point to this discussion. We think that approaching higher-cognition from an ecological basis leads to a fundamentally different flavor of analysis to the typical cognitive approach, one which places less emphasis on representational system features reflecting the influence of computer science on cognitive science and more emphasis on action selection and control. We attempt a brief example of such an analysis below.

A common problem that seems to demand mental representation is the act of talking about something imaginary. This is a complex problem if you treat language as a system of reference; when I say the word ‘unicorn’, to what do I refer, since the referent does not literally exist? If, instead, you treat language as a system for selecting the actions of yourself and others (i.e. if it is a tool; Bickhard, 2009; Everett, 2012) then this problem becomes identical to the problem of using information conventionally to select an action (e.g. the bear and the sign example). Our experiences of using the word ‘unicorn’ dictate the kind of tool that it is and the kind of actions that it can select. When asked to describe a unicorn, a speaker might select the speech actions ‘a horse with a horn’ or something similar. From the perspective of the informational and neural components engaged in action selection, talking about imaginary things is exactly the same kind of process as talking about real things, talking about impossible scenarios, considering multiple possible outcomes, and imagining how things might have been different.

The analysis above is very brief and we agree that ecological psychologists should tackle these tricky problems head on, preferably accompanied by lots of data. But, we hope to have shown that recognizing the role of information in action selection and its relationship to the consequent neural activity does provide the necessary foot in the door (and a necessary vocabulary) for an ecological analysis of higher-order cognition.

## Is This Analysis Representational? Does It Have To Be?

We have relied quite heavily on the notion of designation, which describes when one thing stands in for and is used in place of another thing that you actually want to interact with but cannot, for some reason (Newell, 1980). We argued that this is a useful framing within which to explain what ecological information and the resulting neural activity is for, to an organism.

If this sounds to the reader like we are describing a representation, you would, according to many, be right. Ramsey (2007) has analyzed the concept of cognitive representation in great detail, and emphasizes that while there is little agreement on exactly how cognitive representations work or the format they work in, in order to have any bite, representation has to mean something more than simply an “inner” or “causally relevant” state. (Ramsay, 2007 p 8). Ramsey concludes that regardless of format or other bells and whistles, the essence of all cognitive theories of representation is “the basic idea that inner structures in some way serve to stand for, *designate*, or mean something else” (Ramsay, p 3, emphasis ours).

It may therefore be entirely reasonable to treat both information and the consequent neural activity as representations. These representations are, however, not the computational mental representations of mainstream cognitive psychology, nor could they simply be swapped into such theories. Unlike traditional representations, which must enrich, model or predict sensory data, the psychological power of our decoupled neural activity can *only* comes from its relationship to external ecological information and the property it designates, and that information is given to, rather than invented by, the nervous system (Wilson, 2018a).

Is it worth re-conceptualizing information and the consequent neural activity as representations, though? After all, ecological psychology has always been explicitly anti-representational and a key feature of the current analysis (ecological information solves the poverty of stimulus problem and thus redefines the job description for the brain) has historically been used to motivate non-representational approaches to cognitive science (e.g. Chemero, 2009; also by us, Wilson & Golonka, 2013). While it’s not compulsory to go from designation to representation, there are two reasons why it may be a useful framing to pursue.

First, like it or not, the motivations we identified (especially thinking about thing in their absence) are features of cognition that need to be explained, and perceptual information used for the control of action does not solve the problems by itself. An expanded notion of how information can be used (i.e., selection) and how it works with the nervous system is required. We have shown here that this depends on the issue of designation, so if we want an ecological neuroscience and an account of ‘representation-hungry’ cognition, we quite naturally end up in representational territory. If this ends up being productive, we’ll note that we’ve also shown that information and the consequent neural activity is grounded and can support system-detectable error, so at least they will be good representations.

The second reason is that removing the embargo on the word “representation” may bring the nervous system, its learning history, and its state in a current context back to the main stage. Gibson made a public relations mistake calling his approach ‘direct perception’. The name makes perfect sense, but it has come to imply (for supporters and critics alike) that perception is a free ride; that we simply ‘see’ the world, no cognitive or neural gymnastics required. This isn’t true. Ecological information *does* specify the world and *is* a good stand-in for the world, but it *is not* the same as the world. This means that a) we have to learn to use it as a stand-in (development and perceptual learning should feature front and center of all our theories, e.g. Wicklegren & Bingham, 2001) and b) we will only be able to interact with the world in terms of how it has been projected into information. This last point places meaningful constraints on our search for explanations of our behavior and our neural activity (Golonka & Wilson, 2012, 2018).

This error, of treating the world as simply a given to a cognitive agent, is quite real. In recent years efforts have been made to extend the ecological approach into domains beyond perception and action, especially language and social psychology (Chemero, 2009; Heft, 2007; Kono, 2009; Rietveld & Kiverstein, 2014; Schmidt, 2007) but also the brain (Bruineberg & Rietveld, 2014). As we noted earlier, these efforts all extend Gibson’s notion of affordances to become opportunities for linguistic or social actions that simply account for the behavior of interest. But, affordances, while interesting, are properties of the world, and must be perceived. The critical question (as Gibson himself emphasized) is actually about whether and how these properties are informationally specified, and, despite some efforts (Bruineberg, Chemero & Rietveld, 2018) there is currently no good account for that, nor is one likely (Wilson, 2018b). We have argued recently (Golonka, 2015) that any extension of the ecological approach into so-called higher-order, ‘representation-hungry’ cognition therefore requires extending our understanding of the form, content and use of information, which is what we have continued here. If a broader ecological cognitive science and neuroscience is to become a reality, this is the path we need to take.

All that said, nothing in our analysis hinges on calling neural correlates of ecological information variables ‘representations’. All that matters are the more specific details of the designating nature of information and the related neural activity we have detailed here.

## Summary

A good theory of cognition must account for three key features; implementing intentionality in a physical system, addressing poverty of stimulus concerns to allow intentionality with respect to things currently present, and identifying how we can be intentional with respect to things that are not currently present. Fulfilling this job description led to a strong case for computational mental representations as key players in our theories of cognition, because such systems potentially have the necessary properties to address all three motivations (Fodor, 1980; Newell, 1980; Pylyshyn, 1989). However, these kinds of representations are not a slam-dunk solution. If you have 20 years to spare, we encourage you to peruse this literature and note the many ongoing debates about how to exactly implement these representations, and the various consequences of the methods. When you are done, we hope you agree with our modest assessment that there are certain important concerns with mental representations that have not been laid to rest. In particular, we have discussed that the problems of symbol-grounding and system-detectable error remain largely unsolved.

In this paper we have argued that ecological information and the neural activity it creates can address all three features without falling foul of the problems. Our analysis is grounded in the notion that these components function to designate spatially and temporally distal properties to an organism, and more importantly, it is grounded in the specific way they achieve this designation (via the law-based process of specification). This fact of designation does mean that designating information and neural activity can be considered representational. We are not particularly concerned by this implication, because even if we call these “representations”, the fundamental analysis remains firmly grounded in the ecological ontology. We propose that this framework is best suited for expanding the scope of the ecological approach beyond the real-time control of action and into explaining more abstract cognition and the neural activity that supports it.

## Footnotes

1 But see Picninini, 2004, 2008 for an argument against this

2 Although note that the relationship between mental and neural states (i.e., the neural code) remains a major unsolved issue in cognitive neuroscience.

3 Not everyone thinks the problem remains (Steels, 1997; Taddeo & Floridi, 2007). But the debate remains (Bringsjord, 2015; Wilson, 2018a) and so, for the people who, like us, still worry about this problem, we discuss it here and later, endeavour to solve it.

4 Thanks to Elliott & Sylvia Golonka for inspiring this example

5 The existence of specific kinematic projections provided an evolutionary resource to guide the selection of increasingly sophisticated perceptual systems, which enable behavior to be coupled to and coordinated with distal properties. In fact, we would argue that those seeking to understand why perceptual systems have evolved the form they do should begin with a careful analysis of the spatiotemporal structure of the types of information to which they are sensitive.

6 Selection is also referred to as coordination in the literature.

7 This notion of conventionality has much in common with Pattee’s ideas about symbols in biological systems (e.g., Pattee, 2012).

8 This example is language based, but the idea is effectively classical conditioning, an ability which is ubiquitous in the animal kingdom, so note that ‘conventions’ do not have to be linguistically expressed.

9 Another example is how trained language users experience the heart symbol in the famous ‘I ♥ NY’ as if it were the spoken word “heart.”

10 S*ystematicity* is also discussed, but Johnson (2004) effectively argued against the evidence for and necessity of systematicity in language and thought, so we will remain focused on the other three characteristics here.

11 We mean this as an explicit analogy to EJ Gibson’s (1969) differentiation theory of perceptual learning.

12 The literature on the limits of classical conditioning and chains of associations may be informative for future refinements of this analysis

